# Biofilm Interaction Mapping and Analysis (BIMA): A tool for deconstructing interspecific interactions in co-culture biofilms

**DOI:** 10.1101/2021.08.03.454817

**Authors:** Suzanne M. Kosina, Peter Rademacher, Kelly M. Wetmore, Markus de Raad, Marcin Zemla, Grant M. Zane, Jennifer J. Zulovich, Romy Chakraborty, Benjamin P. Bowen, Judy D. Wall, Manfred Auer, Adam P. Arkin, Adam M. Deutschbauer, Trent R. Northen

## Abstract

Pseudomonas species are ubiquitous in nature and include numerous medically, agriculturally and technologically beneficial strains of which the interspecific interactions are of great interest for biotechnologies. Specifically, co-cultures containing *Pseudomonas stutzeri* have been used for bioremediation, biocontrol, aquaculture management and wastewater denitrification. Furthermore, the use of *P. stutzeri* biofilms, in combination with consortia based approaches, may offer advantages for these processes. Understanding the interspecific interaction within biofilm co-cultures or consortia provides a means for improvement of current technologies. However, the investigation of biofilm based consortia has been limited. We present an adaptable and scalable method for the analysis of macroscopic interactions (colony morphology, inhibition and invasion) between colony forming bacterial strains using an automated printing method followed by analysis of the genes and metabolites involved in the interactions. Using Biofilm Interaction Mapping and Analysis (BIMA), these interactions were investigated between *P. stutzeri* strain RCH2, a denitrifier isolated from chromium (VI) contaminated soil, and thirteen other species of pseudomonas isolated from non-contaminated soil. The metabolites and genes associated with both active co-culture growth and inhibitory growth were investigated using mass spectrometry based metabolomics and mutant fitness profiling of a DNA-barcoded mutant library. One interaction partner, Pseudomonas fluorescens N1B4 was selected for mutant fitness profiling; with this approach four genes of importance were identified and the effects on interactions were evaluated with deletion mutants and metabolomics.

**IMPORTANCE:** The Biofilm Interaction Mapping and Analysis (BIMA) methodology provides a way to rapidly screen for positive and negative interspecific interactions, followed by an analysis of the genes and metabolites that may be involved. Knowledge of these may offer opportunities for engineered strains with improved function in biotechnology systems. *P. stutzeri*, an organism with wide-spread utilization in consortia based biotechnologies, was used to demonstrate the utility of this approach. Where little is known about the factors influencing biofilm based interactions, elucidation of the genes and metabolites involved allows for better control of the system for improved function or yield.

## INTRODUCTION

Consortia based systems in biotechnologies are widespread, however controlling them is challenging due to the genomic and metabolomic complexities of the interactions. Characterization of the genes and metabolites involved in the interactions opens up the possibility for improved consortia functionality by use of engineered strains and culture condition metabolite amendments. Previously, metabolomics of adjacently printed cultures of *P. stutzeri* and *Shewanella oneidensis* were analyzed using replication-exchange-transfer and nanostructure initiator mass spectrometry; however this approach does not elucidate the genes important for the interactions (1). The ability to incorporate genomics into these types of approaches will allow for a better understanding of the interactions involved. Barcoded mutant libraries are proving to be a powerful tool for the discovery of the genetic determinants of co-culture fitness (2, 3). Next-generation sequencing enables rapid profiling of the abundance of barcodes mapped to specific genes in transposon mutant libraries under a wide range of environmental conditions (4, 5) and when integrated with metabolomics, provides rapid functional assignment of transport and metabolic processes (6) important in microbial interactions.

Pseudomonas are a diverse genus of microbes that have been isolated from all over the world of which both beneficial and pathogenic strains have been identified (7, 8). As common soil dwelling microorganisms, they are important constituents of microbial ecosystems and rhizosphere environments (9–12). *Pseudomonas stutzeri*, a model denitrifying pseudomonas, can grow in diverse conditions (13) and its use has been demonstrated in a number of bioremediation processes, including phenol (14), carbon tetrachloride (15), uranium (16), polycyclic aromatic hydrocarbons (17) and diesel oil (18). Additionally, *P. stutzeri* has been discovered and used in a number of consortia based applications, such as bioremediation (19–21), wastewater denitrification (22, 23), aquaculture water quality management (24, 25), as a plant growth promoter for the biocontrol of phytopathogens (26, 27) and manufacturing/municipal waste management (28, 29). The use of *P. stutzeri* biofilms for various bioremediation efforts has potential benefits in terms of activity and yields for copper removal (30), drinking water denitrification (31) and naphthalene (32) and phenol (33) degradation. Given this widespread utilization in both microbial consortia systems and biofilms, *P. stutzeri* was selected to demonstrate the deconstruction of its interactions with other pseudomonas in biofilm-based co-cultures into the genetic and chemical aspects influencing the co-colony fitness.

Here we introduce Biofilm Interaction Mapping and Analysis (BIMA), an integrated platform of automated colony printing, barcoded mutant library profiling and metabolomics to discover and deconstruct the interactions within biofilm-based consortia (Fig. 1). Overlaid colonies were printed using an automated liquid handling system to investigate the morphological, inhibitory and invasive interactions between *P. stutzeri* and other pseudomonas soil isolates in a lab model for a biofilm based consortia. The overlaid colonies were further analyzed using transmission electron microscopy to investigate changes to the species interface over time. Using a DNA-barcoded mutant transposon library of *P. stuzeri* strain RCH2, we identified genes associated with the fitness of the RCH2 in the co-colony. Additionally, exometabolomic analysis was used to evaluate the exchange of metabolites between RCH2 and another pseudomonas strain. We foresee these tools being valuable resources for both the understanding of natural interactions between pseudomonas in microbial communities and in the development of biofilm based biotechnological applications.

**Fig. 1.**
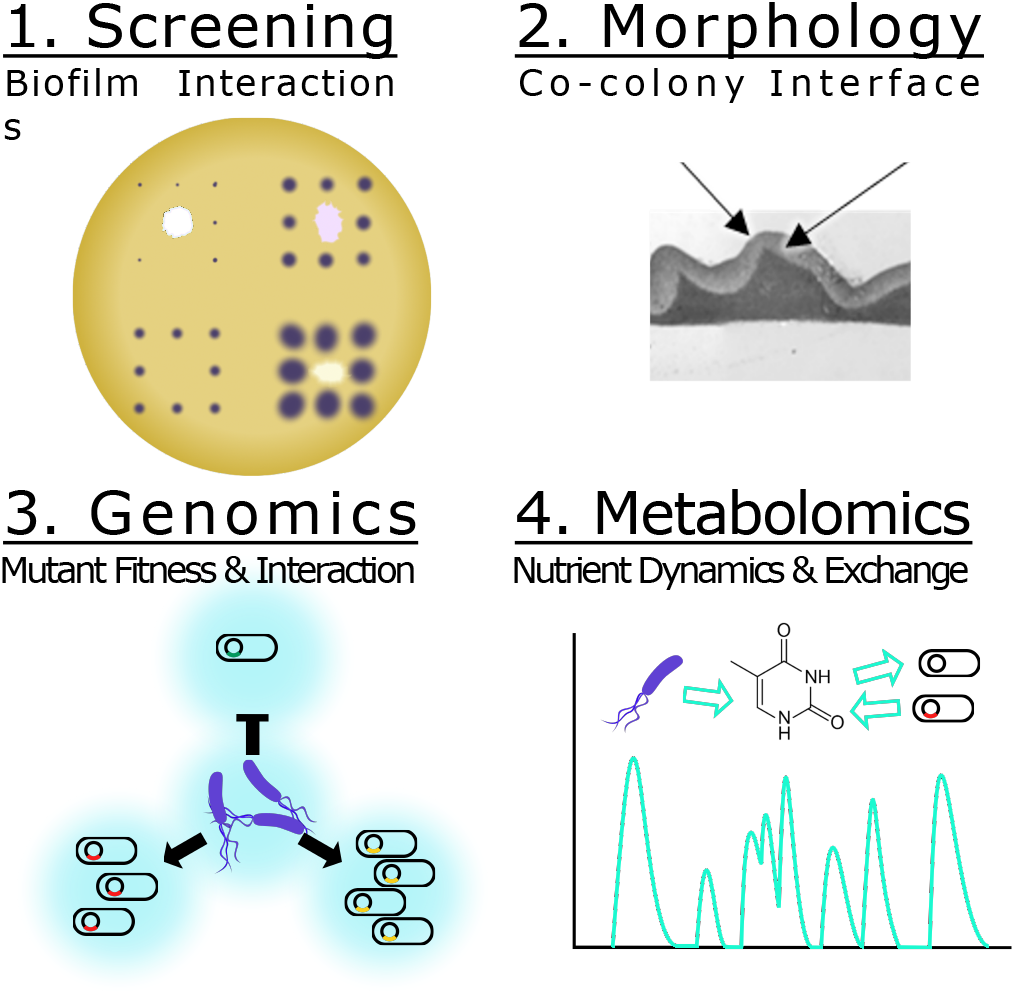
Biofilm Interaction Mapping and Analysis (MIMA) of the biochemical and biophysical interactions required for co-colony fitness. (A) An acoustic printer or automated liquid handling system is used to print underlay colonies of different strains spaced apart to reduce interactions and a single strain grid of overlay colonies to evaluate the macroscopic interactions of the co-colony formation including: inhibition of colonization, direct overlay growth, morphology, color and motility. (B) Transmission electron microscopy is used to evaluate the interface between the two co-colony strains and the changes over time. (C) Mutant fitness profiling of a DNA-barcoded transposon mutant library of the overlay colony allows for investigation of genes important for co-colony formation and fitness. (D) Mass spectrometry based metabolomics is used to analyze the metabolites consumed/released in the co-culture versus mono-culture.

## MATERIALS AND METHODS

### Bacterial strains and growth conditions

*Pseudomonas stutzeri* RCH2 was isolated from chromium(VI) contaminated groundwater at the Department of Energy Hanford 100 Area, Benton County, WA (34). Thirteen additional pseudomonas strains were isolated from Oak Ridge National Laboratory, Field Research Center, TN under conditions indicated in Supplemental table S1. Liquid cultures were inoculated into 3-(N-morpholino)propanesulfonic acid (MOPS) buffered casein yeast magnesium broth (Mb-CYM): 10 g/L pancreatic digest of casein, 5 g/L Difco yeast extract, 1 g/L MgSO_4_ heptahydrate, 10 mM MOPS. Mb-CYM agar was prepared with the addition of 1.5% w/v of agar. Unless otherwise noted, culture conditions were as follows. Strains were maintained as glycerol stocks; revived cultures were plated onto agar plates, incubated at 30°C for 24 hours and stored at 4°C. Starter cultures for experiments were prepared from single colonies inoculated into liquid Mb-CYM broth and cultured aerobically with shaking at 30°C overnight. Uninoculated and unstreaked but incubated cultures/plates were used as negative controls for contamination. In figures and text, *Pseudomonas stutzeri* RCH2 is referred to as culture #1, while the thirteen other strains are referred to as cultures #2-14, and uninoculated controls as culture #15 as indicated in Supplemental Table SI.

### Printed colony biofilm morphology screening assay

Rectangular petri dishes with ANSI standard dimensions were poured with agar to a height of 5 mm. Overnight cultures of the fourteen effector strains (including RCH2) and uninoculated control broth (Supplemental table S1) were diluted 1:50 in fresh Mb-MYM and then added to a 96-well plate. The plates were loaded onto a Hamilton Vantage Liquid Handling System equipped with a 96 pipetting head which was used to dip pipette tips into the liquid cultures and press/print them against the surface of the agar in a grid; a single colony of each unique strain was printed in a 3 × 5 grid (Fig. 2A). The lidded-plate was sealed with Parafilm and incubated at 30°C for two days once visible colonies had formed of similar size (Supplemental figure S1). Printing was repeated using only *P. stutzeri* RCH2 (the interaction strain) in all well positions of a 16 × 24 grid (ANSI standard dimensions of a 384 well plate), excluding the positions from the original print at time 0 (Fig. 2A). Cultures were incubated for an additional 4 days prior to imaging using a digital camera. Image analysis was performed using ImageJ (35). Briefly, the image was converted to 16-bit grayscale, corrected for uneven lighting using the FFT bandpass filter (filter large structures to 70 pixels and small structures to 5 pixels, 5% tolerance, autoscale and saturate after filtering), auto thresholded to black and white, and then the colonies were measured using the Analyze Particles tool (colonies missed by the automated selection were manually outlined and then measured). Measurements included area and position. Colony areas of surrounding (8 colonies in a square around the center interaction colony) and closest/farthest neighbors (sides/corners of the square) were compared with the colonies surrounding the control uninoculated position (#15) using Dunnett’s multiple comparison procedure using R version 3.6.2 (36, 37).

**Figure 2.**
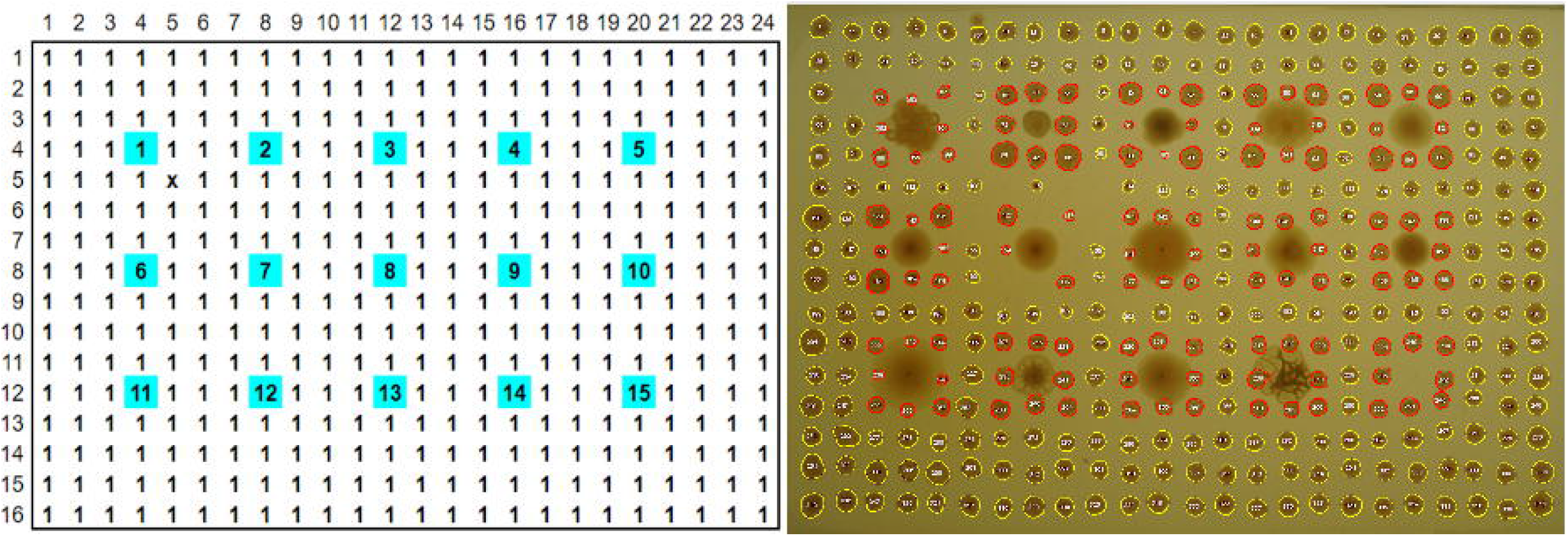
Colony interaction screening. (A) Fourteen Pseudomonas “effector” strains (spots #1-14) and one uninoculated control medium (spot #15) were printed from liquid cultures onto an agar plate (blue). After 2 days of incubation, the interaction strain, *P. stutzeri* RCH2 (#1) was printed in all other positions (white). (B) Image taken at 6 days total incubation time indicated inhibited growth of *P. stutzeri* RCH2 around strains 1 (self), 3, 4, 5, 6, 7, 8, 9 and 11. (C and D) *P. stutzeri* RCH2, the interaction strain, colony size was used to evaluate which effector strains had a significant (ANOVA, Dunnett’s test, p<0.001 ***, p<0.01 **, p< 0.05 *) effect on the growth of the neighboring RCH2 colonies. One effector strain (#2), N1B4, with the largest mean colony area and not inhibited by RCH2, was selected for further morphological and metabolic interaction with RCH2. When comparing all surrounding colonies, only 7 was significantly different from control (Supplemental fig. S2).

### Co-colony overlay morphology with light microscopy and transmission electron microscopy

Overnight culture of *Pseudomonas* sp. FW300-N1B4 (N1B4) was manually pipetted (10 uL) onto an agar plate and incubated at 30°C overnight. Overnight culture of *P. stutzeri* RCH2 was manually pipetted (10 uL) over the top of the N1B4 colony with care to not puncture the colony surface and to avoid allowing the droplet to spill over the sides. For time course analysis of macroscopic morphology, the rim of the petri dish was sealed with parafilm and placed on a black velvet cloth under a Leica M165FC microscope with a Planapo 1.0 × objective lens. Images were acquired every 2 minutes for a 24 hour period with a Leica DFC400 1.4 megapixel ccd sensor digital camera set at a 9 mm frame width. For microscopic colony interface analysis, after overnight growth of the overlay, the colony and underlying agar was cut out from the plate and fixed for 2 hours with 2.5% glutaraldehyde in 0.1 M sodium cacodylate buffer (pH 7.2). Fixed samples were stained with 5 mM ruthenium red in 0.1 M sodium cacodylate buffer (pH 7.2) for 1 hour and post-fixed with 1% osmium tetroxide for 1 hour. Following staining, samples were dehydrated through a graded ethanol series (20%, 40%, 60%, 80%, 90%, 100%, 100%, 100%) followed by infiltration with an Embed-812 epoxy resin (Electron Microscopy Sciences):acetone series of 1:3 for 2 hours, 2:3 for 2 hours, and 100% resin overnight. Samples were then heat polymerized in 100% resin with N,N-dimethylbenzylamine accelerant for 2 hours at 85°C. 100 nm sections were cut and sectioned using a Leica UC6 Ultra Microtome (Leica Microsystems Inc., Buffalo Grove, IL, USA) and then stained sequentially with 2% methanolic uranyl acetate and Reynolds’ lead citrate for 5 minutes each. Images were collected using a FEI Tecnai 12 transmission electron microscope (FEI Company, Hillsboro, OR, USA).

### Fitness profiling of RCH2 mutants in the presence of other pseudomonads

The construction of the *P. stutzeri* RCH2 DNA-barcoded transposon mutant was previously described (4). Overnight culture of *Pseudomonas* sp. FW300-N1B4 was manually pipetted (10 uL) in triplicate onto LB agar plates. The plates were incubated overnight and then 5 uL of the *P. stutzeri* RCH2 barcoded transposon mutant library (in LB media) starter culture at an OD_600_ of 0.9 was spotted as overlay colonies on top of the three replicates of the underlay colonies. An aliquot of the starter culture was used for the initial RCH2 mutant abundances. Colonies were then carefully scraped and frozen for storage and subsequent analysis of the final *P. stutzeri* RCH2 mutant abundances. Gene fitness scores were calculated by comparing the initial and final mutant abundances as determined by deep sequencing of the DNA barcodes, as previously described (4).

### Mutant construction

The four mutants from the pooled fitness assay with the largest absolute differential fitness where fitness of RCH2 with N1B4 was less than fitness of RCH2 on agar alone were selected for constructing gene deletion mutant strains. Deletion mutants (Supplemental table S2) for gamma-glutamyl phosphate reductase, OHCU decarboxylase, formyltetrahydrofolate deformylase and glutamate 5-kinase were constructed by conjugation of unstable, marker-exchange plasmids into *P. stutzeri* RCH2, as previously described (38) with modifications as follows. All plasmids and primers (IDT, Newark, NJ) used are listed in Supplemental tables S3 and S4, respectively. Briefly, deletion cassettes, containing kanamycin resistance gene (npt) flanked by chromosomal regions up and downstream of the gene to be deleted, were assembled using the “gene SOEing” technique (39). Cloned homologous regions were sequenced at the DNA core facility at the University of Missouri, Columbia, and compared with the published sequence for *P. stutzeri* RCH2 (40). Cassettes and the template plasmid pM07704 were amplified by polymerase chain reaction (PCR) with Herculase II DNA polymerase (Stratagene). Marker-exchange plasmids were generated by ligation of the PCR products in α-select cells (Bioline) using the SLIC cloning method (41). The plasmids were isolated and transformed into *E. coli* strain WM3064, and then transferred to RCH2 via conjugation (42). The plasmids, containing up/downstream chromosomal regions flanking npt, allow for exchange of npt with the gene of interest in RCH2 via double homologous recombination. Exconjugates were selected on 50 ug/mL kanamycin solid medium, then screened for spectinomycin (100ug/mL) sensitivity to ensure no single recombination isolates were selected. Deletion strains were confirmed by Southern blot analysis. One isolate for each deletion of interest was retained while the other isolates were discarded. All strains were frozen as early stationary phase cultures in 10% (v/v) glycerol. RCH2 mutants used for experiments were maintained in Mb-CYM broth or Mb-CYM agar with 50 ug/mL kanamycin sulfate.

### Mutant co-colony morphology analysis

Overnight cultures of RCH2 (wild-type and mutants) and N1B4 (500uL) were centrifuged (5000xg × 3min at RT) to collect cells, then washed 2 times in DPBS (resuspension in 500uL DPBS, centrifugation at 5000xg × 3 min at RT) with final resuspension adjusted to an OD (600nm, 1cm) of 0.5. Uninoculated control medium was washed and resuspended in a similar manner to account for carryover of nutrients from tube surfaces and residual volumes. Cultures (“underlays”) were manually spotted (2 uL) onto Mb-CYM agar plates in a 7 × 7 format and incubated overnight at 30°C. Fresh cultures were inoculated from agar stock plates into Mb-CYM. Overlay colonies were diluted and washed in the same manner as for the underlays and then carefully spotted on top of the underlay colonies. Images were taken using a digital camera of whole plates and through a LEICA M165 FC stereo microscope (2x objective) at 24 hours (0 hours of overlays), 48 hours (24 hours of overlays), 72 hours (48 hours of overlays). Morphological differences were visually evaluated.

### Metabolomics analysis

Wild-type and mutant strains of *P. stutzeri* RCH2 were cultured in ‘spent’ N1B4 medium (sterile filtered N1B4 culture supernatant) in liquid culture and then exometabolites were collected for LCMS analysis. For collection of ‘spent’ medium, overnight liquid cultures of RCH2 wild-type, RCH2 mutants, and N1B4 wild-type and uninoculated control medium were centrifuged (3000 × g × 10 minutes) to pellet cells. Supernatants were sterile filtered (0.22 um), supplemented with 1x Wolfe’s vitamins and minerals (ATCC) and stored at 4°C overnight. Additional overnight cultures were diluted, washed and adjusted as described above for mutant co-colony morphology analysis, except the final OD was adjusted to 0.12 in DPBS, 20 uL of which was added to 100 uL of spent medium in 96 well plates. Cultures were prepared in triplicates for all combinations of mutants and wild-type RCH2 on spent N1B4 and fresh Mb-CYM and for N1B4 on all combinations of spent mutants and wild-type RCH2 and fresh Mb-CYM; uninoculated controls were included for each spent medium and positive growth controls were included for each strain on fresh (non-spent) medium. Cultures were incubated for 20 hours in a BioTek plate reader with OD readings taken every 30 minutes at 600nm. Cross-feeding medium from the culture was then collected via centrifugation (3000x g for 5 minutes), transferred to a new plate that was sealed with heated foil (4titude), and frozen at -80°C. Holes were pierced in the foil with an 18 gauge needle and then supernatants were lyophilized to dryness. Dried material was resuspended in 200 uL of internal standard mix (15uM 13C, 15N amino acid mix, 10 ug/mL 13C-mannitol, 13C-trehalose and 2 uM 15N4-inosine, 15N5-adenine, 13C4-15N2-uracil, 15N4-hypoxanthine, 13C4-15N2-thymine) in LCMS grade methanol, resealed, vortexed, sonicated in a room temperature water bath for 10 minutes, rechilled at - 80°C for 5 minutes, and centrifuged 3000x g for 5 minutes to pellet insoluble material. Supernatants were filtered (0.22µm, PVDF) using an Apricot positive pressure filtration device. Filtrates were then arrayed into 50uL aliquots in a 384 well plate for LCMS analysis.

### LCMS analysis of exometabolites from spent media fed cultures

Extracts of polar metabolites were analyzed using hydrophilic interaction chromatography - mass spectrometry. Metabolites were retained and separated on an InfinityLab Poroshell 120 HILIC-Z column (Agilent, 683775-924, 2.7µm, 150 × 2.1 mm) using an Agilent 1290 UHPLC. Samples, held at 4°C, were injected at 4uL each; the column temperature was held at 40°C and flow rate was held at a constant 0.45mL/min. Following injection, a gradient of mobile phase A (5mM ammonium acetate, 0.2% acetic acid, 5uM methylene di-phosphonic acid in water) and mobile phase B (5mM ammonium acetate, 0.2% acetic acid in 95:5 (v/v) acetonitrile:water) was applied as follows: initial equilibration at 100% B for 1.0 minutes, linear decrease to 89% B over 10 minutes, linear decrease to 70% B over 4.75 min, linear decrease to 20% B over 0.5 min, hold at 20% B for 2.25 min, linear increase to 100% B over 0.1 min, re-equilibration at 100% B for 2.4 min. Eluted metabolites were subjected to mass spectrometry analysis on a Q Exactive Hybrid Quadrupole-Orbitrap Mass Spectrometer (Thermo Scientific) equipped with a HESI-II source probe using Full MS with Data Dependent tandem MS. Source settings were as follows: sheath gas at 55 (arbitrary units), aux gas flow at 20, sweep gas at 2, spray voltage at 3 |kV| spray, capillary temperature at 400C, and S-lens RF at 50. MS1 was set at 70,000 mass resolution, with automatic gain control target at 3.0E06 with a maximum allowed injection time of 100 milliseconds, at a 70-1050 m/z scan range. dd-MS2 was set at 17,500 mass resolution with automatic gain control target at 1.0E5 and a maximum allowed injection time of 50 milliseconds, a 2 m/z isolation window and stepped normalized collision energies at 10, 20 and 30 (dimensionless units). All data was collected in centroid mode. The scan cycle included a single MS1 scan followed by sequential MS/MS of the top two most intense MS1 ions excluding any fragmented within the previous 10 seconds. Ions selected for fragmentation must meet a minimum AGC target threshold of 1.0E3 with absolute charge less than four. Each sample was analyzed in negative ionization mode. Sample were injected in randomized order with solvent blank injections between each; internal and external standards were used for quality control purposes and for retention time predictions of compounds from in an in-house standards library. Using custom python scripts and metabolite atlases (43, 44), mass-to-charge ratios, retention times and where possible spectra fragmentation patterns were used to confirm metabolite identification by comparison to metabolite standards analyzed using the same LC-MS/MS methods. Mutant and RCH2 cultures on N1B4 medium were compared with uninoculated control N1B4 spent medium using ANOVA and Tukey HSD in R.

### Data Availability

Raw LCMS data are available from the JGI Genome Portal (genome.jgi.doe.gov) under project number 1278333.

## Results

### BIMA screening: Microbial interaction mapping and strain selection

As the first step of BIMA, a colony printing method was developed on an automated liquid handling system for the purpose of scalability and transferability between labs. Fourteen preprinted ‘effector’ Pseudomonas strains (including RCH2) were evaluated for their effect on the growth of subsequently printed neighboring colonies of *P. stutzeri* RCH2 (Fig. 2A and 2B). Pseudomonas strains 3-9, and 11 inhibited the growth of the four closest RCH2 colonies, located along the sides of the square of neighboring colonies (Fig. 2C). Interestingly, for a subset of these (3, 4, 5), the more distant RCH2 colonies at the corners of the square of neighboring colonies had larger mean areas (Fig. 2D). Pseudomonas strain 2 (*Pseudomonas fluorescens* FW300-N1B4) was the only strain that did not inhibit RCH2 (closest side colonies) and enhanced the growth of RCH2 (corner colonies). This strain ‘N1B4’, was selected for further analysis in co-culture with *P. stutzeri* RCH2.

#### BIMA morphology: Co-colony structure and infiltration

Time lapse images of the RCH2 overlay on N1B4 were taken to evaluate the co-colony morphology. Highly wrinkled, rugose colony morphology had formed by 12 hours after application of the RCH2 overlay (Fig. 3A). The co-colony species interface was further examined using transmission electron microscopy (TEM) (Fig. 3B). TEM after 2 hours showed a stratification with a clear distinction between RCH2 on the surface and N1B4 underneath. Over the course of 24 hours, the colonies became more mixed with infiltration of the N1B4 layer by RCH2. In both the co-culture and in isolate culture, RCH2 biofilm developed exopolysaccharide sacs containing groups of spherical cells at the air interface; whereas when spotted on the surface of N1B4, single, elongated RCH2 are observed at the N1B4 interface. The sacs became visible in the 24 hour co-colony images and are similar to those observed in other *P. stutzeri* strains where the sacs are associated with oxygen exclusion for nitrogen fixation under aerobic conditions (45). The anaerobic co-colony culture developed a compact RCH2 layer with no visible EPS sacs.

**Figure. 3.**
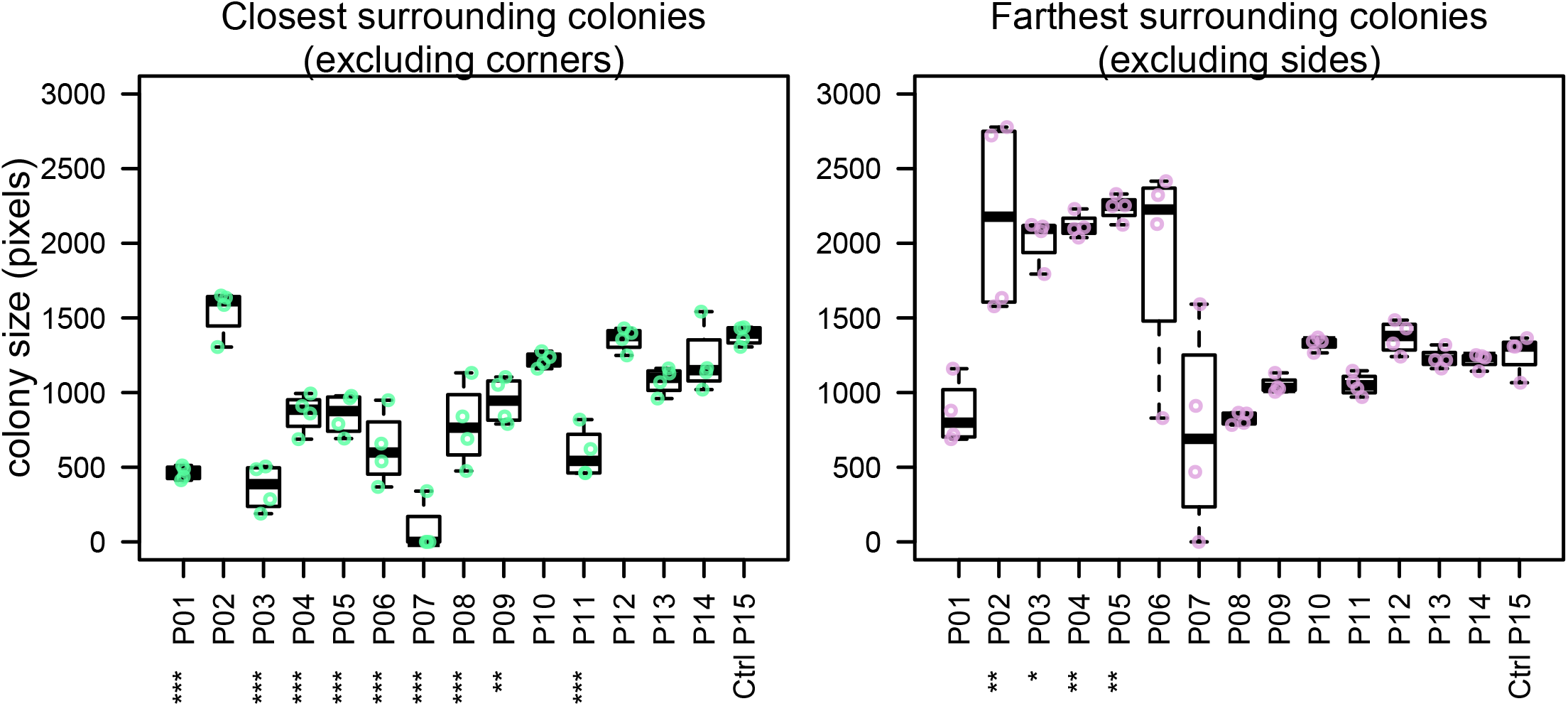
Transmission electron microscopy of co-colony morphology. (A) Timepoint images of a co-colony of *Pseudomonas* sp. FW300-N1B4 (underlay) and *Pseudomonas stutzeri* RCH2 (overlay) were taken every 6 hours for the first 24 hours of the overlay culture. Control N1B4 cultures remain smooth (see Fig. 4 and S02). (B) Transmission electron microscopy imaging (1400x) of vertical slices showing the co-colony structure, air-colony interface, and strain interface were taken at 2, 12 and 24 hours of aerobic RCH2 overlay culture, and at 24 hours of an anaerobic RCH2 overlay culture. Similar imaging at 24 hours of aerobic RCH2 and N1B4 monocultures were used to determine staining and morphology of each cell type.

### BIMA genomics: Mutant fitness analysis and interactions

A pooled mutant fitness assay was used to identify RCH2 genes essential for successful co-culture growth with N1B4. The top four genes with the largest differential gene fitness between RCH2 on LB and RCH2 on N1B4, where fitness on N1B4 was negative included gamma-glutamyl phosphate reductase (*proA*), OHCU decarboxylase (*uraD*), formyltetrahydrofolate deformylase (*purU*), and glutamate 5-kinase (*proB*) (Fig 4A). To investigate these genes further, we constructed isogenic single-gene deletion strains for all four genes.

**Figure 4.**
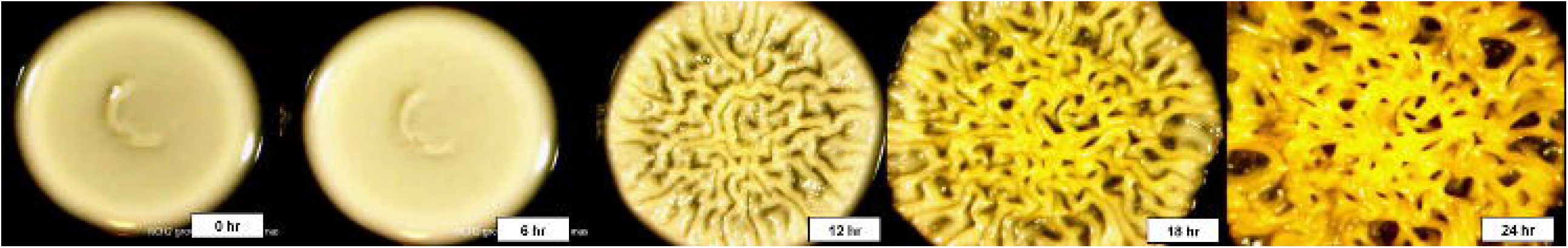
RCH2 mutant co-colony interactions. (A) RB-TnSeq mutant library fitness assay, (B) mutant growth on N1B4 spent and fresh media, (C) mutant growth on N1B4 colonies. All strains of RCH2 grew alone (on control sterile medium underlay) and formed rugose colonies by 24 hours of overlay culture (Supplemental Fig. S3, Supplemental Table S5). Images are notated with “S” or “R” for smooth or rugose morphology, respectively; location noted when rugose morphology was located only in the center or periphery of the colony.

In liquid culture, while all mutants were capable of growth on fresh medium, only the *ΔuraD* deletion strain had similar growth to the wild-type when cultured on spent N1B4 medium (Fig. 4B). Minimal to no growth was observed for the *ΔproA* and *ΔproB* deletion mutants, and delayed growth was observed for the *ΔpurU* deletion mutant. N1B4 had similar growth on the spent medium of RCH2 wild-type and mutants (not shown). The limited or delayed growth indicated these mutants were unable to acquire some required metabolite from the medium that was consumed by N1B4 in liquid culture or that growth may have been inhibited by something N1B4 was producing.

The biofilm interactions were analyzed by manual printing of the RCH2 wild-type and mutant strains on the top of an established N1B4 colony. The overlays were imaged at 24 hours and 48 hours after overlay printing (Fig. 4C). Wild-type RCH2, and *ΔproA* and *ΔuraD* deletion strains formed rugose co-colony biofilms after 24 hours similar to the wild-type but with a smoother center for *ΔproA*, while the *ΔpurU* mutant had delayed rugose formation and the *ΔproB* mutant remained smooth up to 72 hours of observation. All mutants formed rugose colonies after 24 hours of overlay on control medium. On its own, the *ΔpurD* formed more tubular structures than the wild-type alone, similar to growth on N1B4 for both (Supplemental Fig. S3).

### BIMA metabolomics: Potential for metabolite exchange

To further evaluate why the four RCH2 genes identified in the mutant fitness assay were important to growth in the co-colony, spent medium from N1B4 was fed to each of the mutants and the wild-type RCH2 and then metabolomics was performed to check for altered metabolism in the mutants. Despite limited growth of three of the mutants on N1B4 spent medium (Fig. 4B), all four RCH2 mutants appeared to be metabolically active with significant increases and decreases in metabolite abundances relative to uninoculated N1B4 spent medium and in many cases were significantly different than the wild-type (Fig 5 and Supplemental Fig. S4). Metabolites in pathways associated with the mutated genes, including one-carbon (C1) metabolism (PurU), ureide pathway (UraD) and proline synthesis (ProA/B) were evaluated further (Supplemental Fig. S05). The *P. stutzeri* RCH2 wildtype consumed several N1B4 metabolites involved in one-carbon (C1) metabolism though consumption of methionine was reduced in *P. stutzeri RCH2* strain JWST9066 (*ΔpurU*) (Supplemental Fig. S05). *P. stutzeri* RCH2 strain JWST9063 (*ΔuraD*) accumulated allantoin, presumably from spontaneous conversion of 2-oxo-4-hydroxy-4-carboxy-5-ureidoimidazoline (OHCU) to (R)-allantoin, which RCH2 would be unable to use in downstream nitrogen assimilation pathways without a racemase (46), an enzyme *P. stutzeri* has been shown to lack (47). *P. stutzeri* RCH2 strain JWST9060 (*ΔproA*) and *P. stutzeri* RCH2 strain JWST9069 (*ΔproB*), both known auxotrophs for proline (48), had reduced consumption of glutamate (precursor in proline synthesis) from N1B4, when compared to wild-type (Supplemental Fig. S05).

**Fig. 5.**
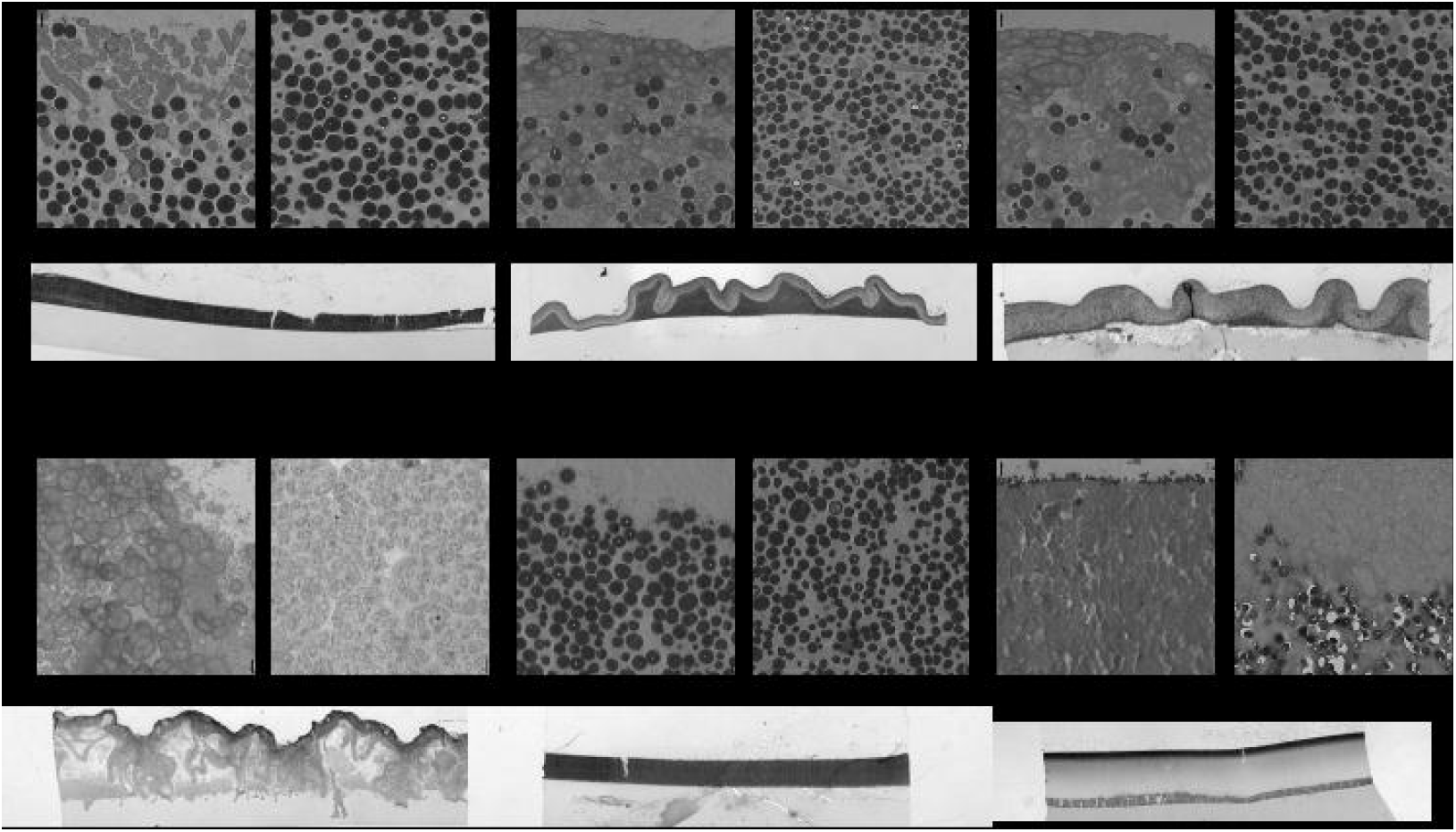
Metabolomics. N1B4 spent medium was sterile filtered and used to culture *Pseudomonas stutzeri* RCH2 wild-type, and mutants *ΔuraD, ΔpurU, ΔproA, ΔproB* for 24 hours before collection of extracellular metabolites for extraction and LCMS analysis. A media control was incubated alongside the sample cultures to control for contamination and for comparison with starting metabolite abundances. A targeted analysis was performed using a library of m/z, retention time and MS2 spectra of common polar metabolites. Metabolites with significantly different abundances relative to control media (ANOVA, Tukey HSD, p<0.05) are indicated with an asterisk. Metabolites with significantly different abundance between at least two of the cultures are indicated with #. Boxplots and statistical comparisons are available in Supplemental Figure S4, S5 and Table S6.

## DISCUSSION

Biofilm formation and morphology in pseudomonas appears to be regulated by a combination of biotic factors including, but not limited to: phenazines, exopolysaccharide production, signaling molecules and flagellar activity (49). In pigmented species of pseudomonas, it has been shown that a reduced cytoplasm (induced under anoxic and low nitrate conditions) as measured by NADH:NAD+ ratio results in a rugose or wrinkled morphology while under aerobic and normal nitrate conditions, where reduction of phenazines or nitrate to dinitrogen gas occurs, they maintain a smooth morphology (50). Interestingly, *P. stutzeri*, has a distinguishing morphological characteristic of forming rugose or wrinkled colonies following initial isolation and culture on agar. While brown in color due to cytochrome c, they are classified as a nonpigmented, nonfluorescent pseudomonas and are not known to produce phenazines(13). *P. stutzeri* can grow anaerobically in the presence of nitrate and has been used for denitrification purposes(13). However, it has been demonstrated that after repeated culture, smooth colonies can form and it may take several transfers in nitrate media under semi-aerobic conditions before they are able to grow under denitrifying anaerobic conditions (13).

Four genes important for survival of *P. stutzeri RCH2* on the surface of *P. fluorescens* FW300-N1B4 were evaluated for their effects on utilization of metabolites from N1B4. From metabolomics analysis, we have postulated possible metabolic interactions that may support the growth of RCH2. Given that PurU is important in maintenance of one carbon pools, the *ΔpurU* mutant may be unable to obtain the reaction products (THF and formate) from N1B4, and as suggested by previous studies in *E. coli*, may experience glycine starvation due to GlyA inhibition by methionine and adenine (metabolites present in N1B4 spent medium) (51). In some bacterial species, the ureide pathway is utilized for recovery of nitrogen from purines under stress conditions(52, 53). Under aerobic conditions or presence of ammonia, N2 fixation in *P. stutzeri* is suppressed and nitrification is active, however in biofilm, nitrogen fixation is suspected to occur in the EPS sacs produced by the rugose morphology(45, 54). At the N1B4 – RCH2 interface of the co-colony (which lacked EPS sacs in imaging) and in liquid culture with shaking and sufficient gas exchange, nitrogen fixation may be suppressed in favor of utilization of exchanged organic nitrogen compounds / intermediate allantoin precursors (5-hydroxyisouric acid or OHCU) from N1B4 spend medium, prior to conversion to usable (S)-allantoin. The wild-type may use exchanged glutamate for proline synthesis while the proline synthesis mutants are likely unable to obtain enough proline directly from N1B4 and is likely unable to sufficiently utilize alternative sources such peptide degradation or by alternative synthesis from ornithine (Supplemental Fig. S05). Together, these results indicate the growth of the RCH2 may be reliant upon uptake of metabolites produced by the N1B4.

While our experiments focused on cooperative interactions, inhibitory interactions may be of interest in controlling pathogenic strains in agricultural systems. The inhibitory interactions of certain pseudomonas species have been studied in detail. For example, some pathogenic species of pseudomonads produce lipodepsipeptides (e.g.. corpeptins from *P. corrugata*, a plant pathogen) which also have antimicrobial activity (55). These may contribute to interspecific competition between pseudomonads by inhibiting biofilm formation and breaking down existing biofilm (56). An interesting follow up study involving a mutant library of *Pseudomonas corrugate N2F2*, may elucidate the mechanism of inhibition between the two species and the nature of its competitive and pathogenic behavior in nature. Additionally, some beneficial strains such as *P. aureofaciens*, an anti-fungal symbiont of wheat, releases phenazines in the rhizosphere which inhibit the growth of plant pathogens in response to exogenously diffusible signaling molecules (57). These compounds may contribute to competitiveness and the ability to form biofilms (58). Interestingly, phenazines may have multiple functional roles (59), including involvement of the redox state of the cytoplasm (50, 60); interactions such as these may be important determinants of co-colony biofilm structure under varying oxygenic conditions.

In this work we have demonstrated that bacterial printing can be used to rapidly screen for macroscopic interactions such as changes to colony morphology, size and color. We take advantage of a mutant fitness library and metabolomics to gain insights into genes and compounds mediating observed bacterial interactions. We anticipate that this approach integrating bacterial printing, mutant fitness libraries, targeted genetics, and metabolomics is suitable to investigating diverse microbial interactions. In some instances cooperative growth may be favorable (such as for bioremediation purposes where both consortia and biofilm based systems are used). In other cases, competitive interactions may be desired for biocontrol based purposes or for understanding competitive interactions in soil environments. Knowledge of the genomic and metabolic determinants involved in these interactions allows for more directed design of co-colony based systems and a better understanding of those existing in nature. We foresee BIMA becoming a valuable tool for the enhancement and understanding of *P. stutzeri* and other co-colony based systems.

## Acknowledgements

Study design, sample collection, data analysis and interpretation, and manuscript preparation by ENIGMA (Ecosystems and Networks Integrated with Genes and Molecular Assemblies, http://enigma.lbl.gov, a Scientific Focus Area Program at Lawrence Berkeley National Laboratory), is supported by the U.S. Department of Energy, Office of Science, Office of Biological & Environmental Research, Genomic Sciences Program under contract number DE-AC02-05CH11231 to Lawrence Berkeley National Laboratory. Data analysis used resources of the National Energy Research Scientific Computing Center, a Department of Energy Office of Science User Facility operated under contract number DE-AC02-05CH11231. The funders had no role in study design, data collection and interpretation, or the decision to submit the work for publication.

---

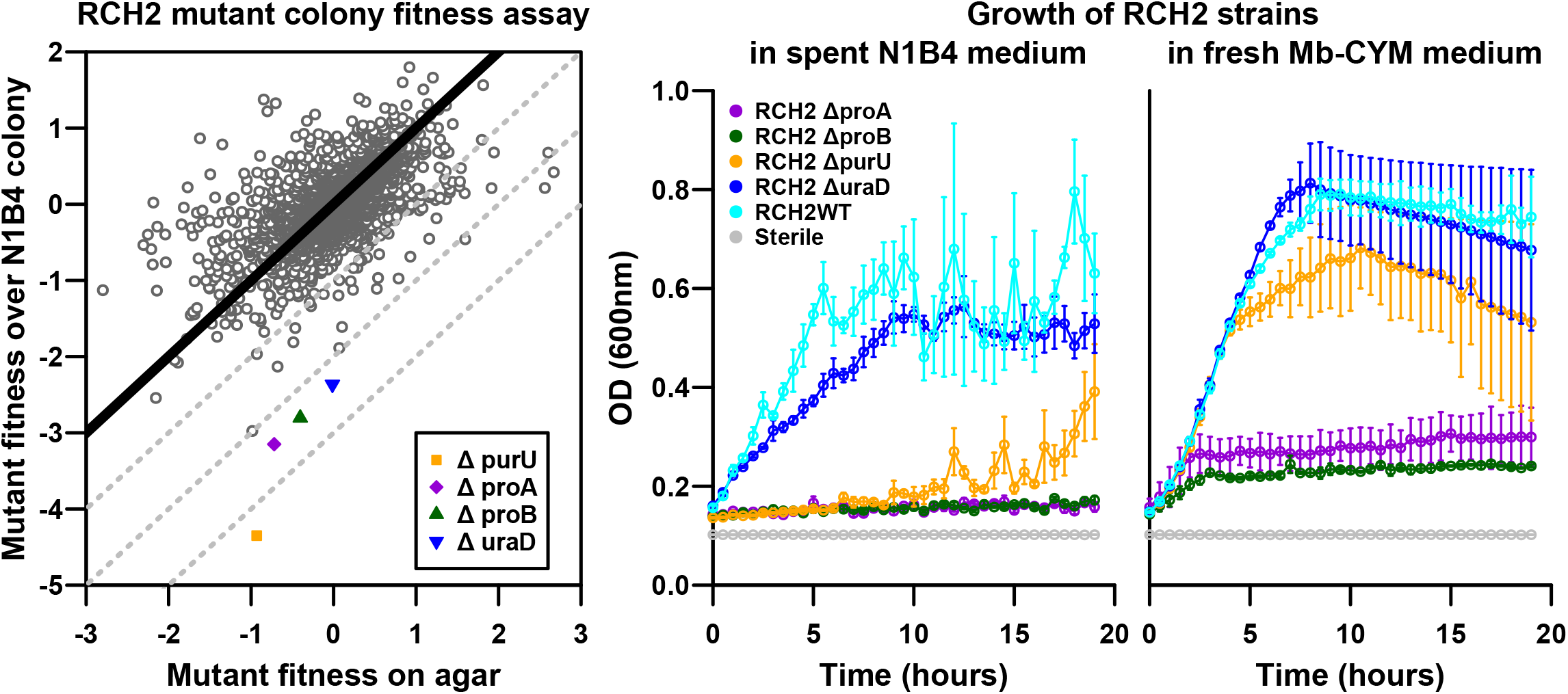

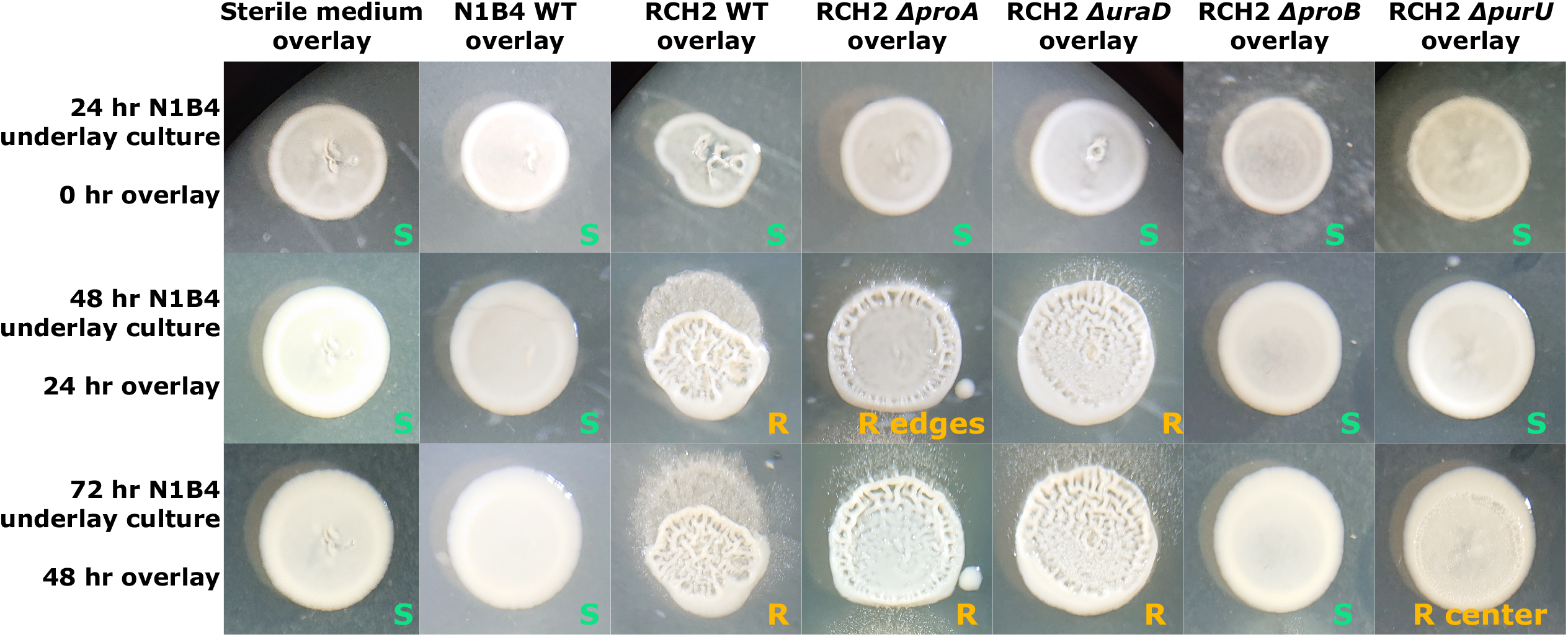

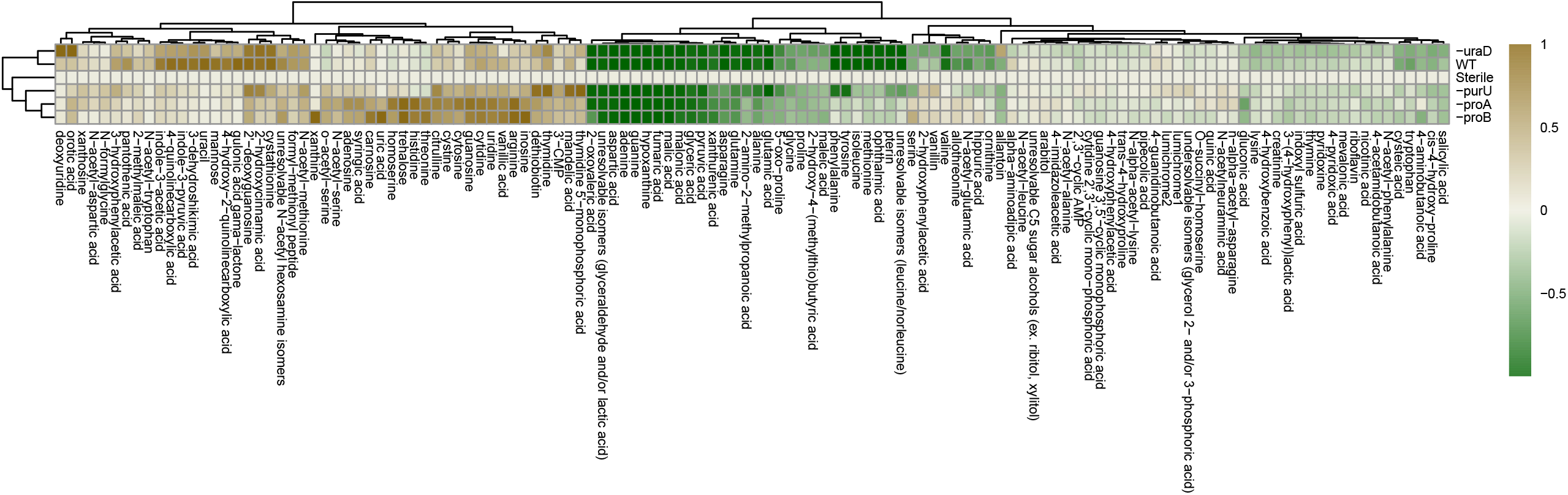

## Notes

### Competing Interest Statement

The authors have declared no competing interest.

