## Supplemental Figures for "Biofilm Interaction Mapping and Analysis (BIMA): A tool for deconstructing interspecific interactions in co-culture biofilms"

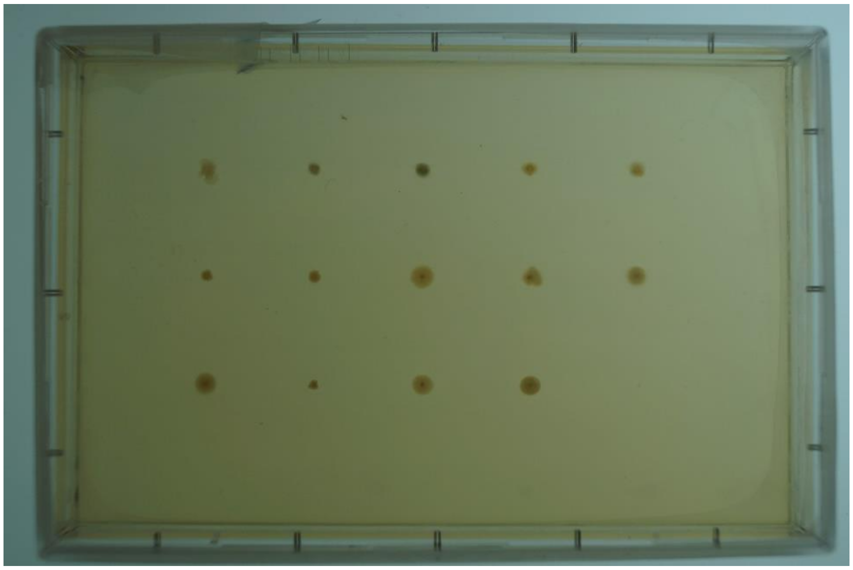

**Figure S01. Morphology screening assay, day 2.** Effector colonies (3x5 grid at 2 days, just before overlay printing. Overlays were printed in a 16x24 grid (excluding positions from the 3x5 grid) using a liquid handling system.

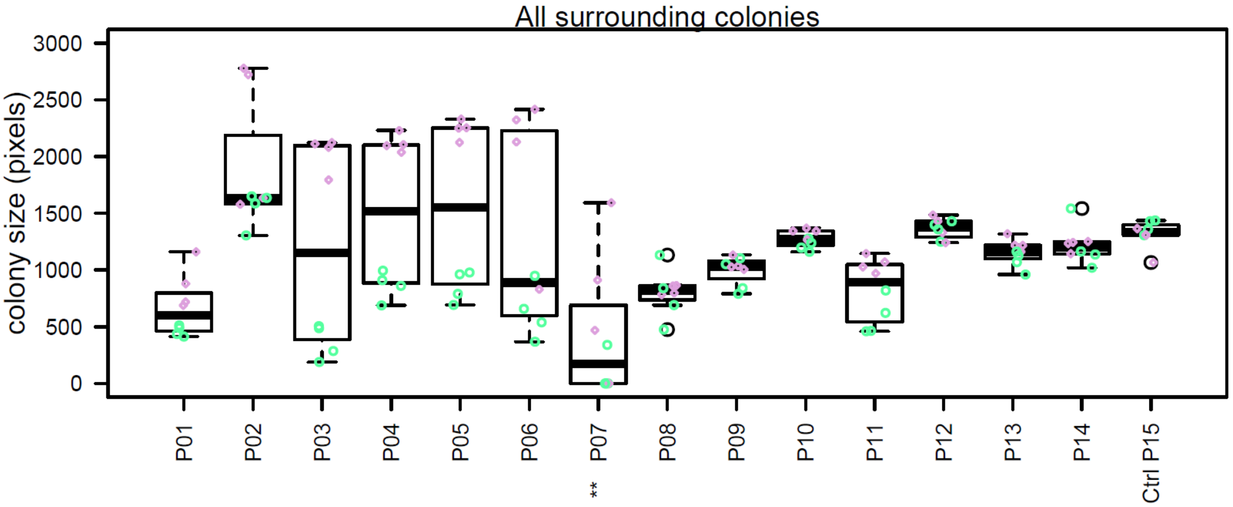


**Figure S02. Colony interaction screening, all surrounding colonies** When comparing all eight colonies of *P. stutzeri RCH2* surrounding each effector strain, including corners (green) and sides (purple), only

culture #7 significantly influenced growth of the interaction strain compared with the control. ANOVA, Dunnett's test, p<0.001 ***, p<0.01 **, p< 0.05 *.

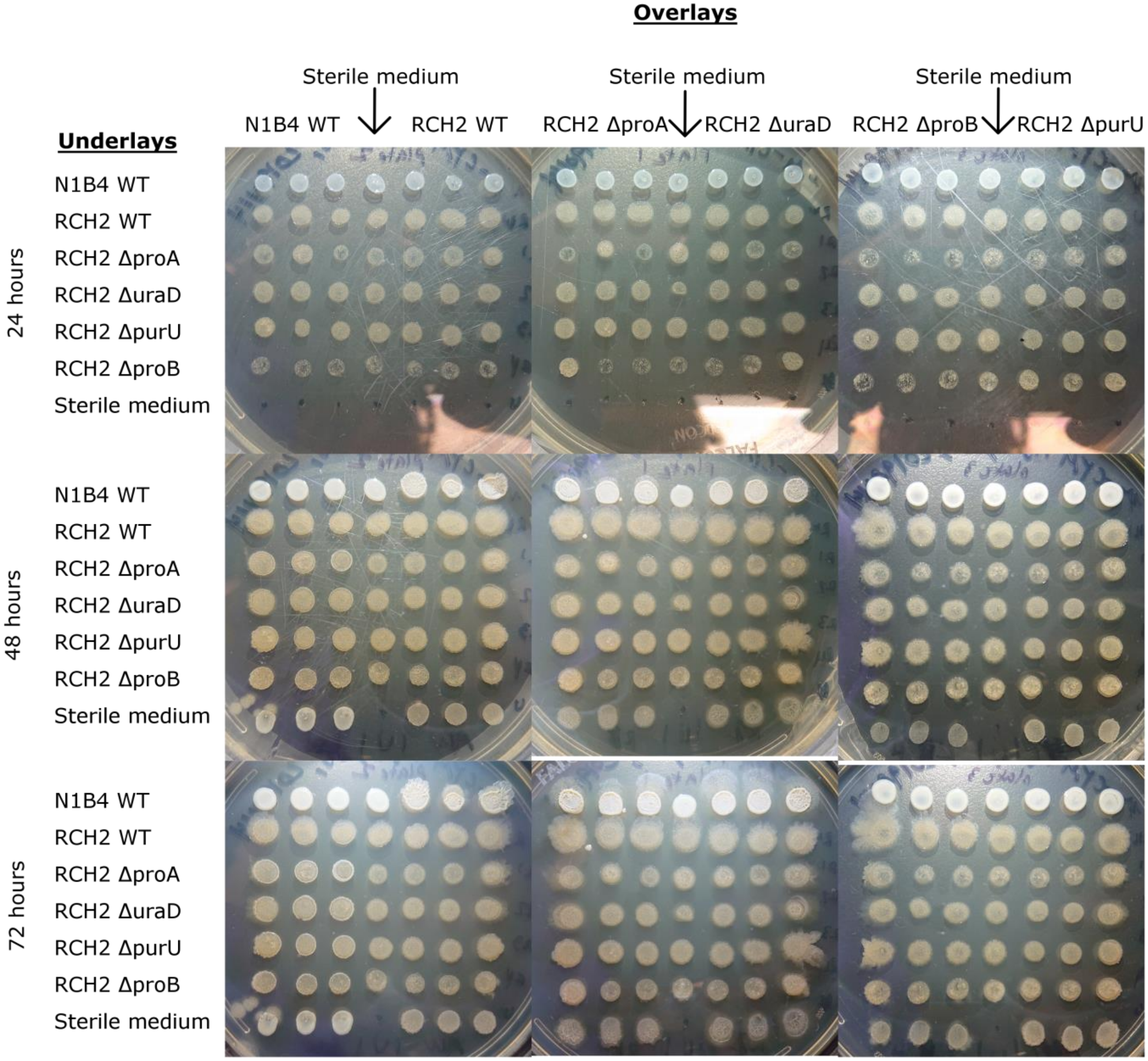


**Fig. S03. Co-colony morphologies of all underlay and overlay colonies at 24, 48 and 72 hours.** Subset of data was used for close up images shown in figure 04. Three replicates of each co-colony were spotted horizontally next to one another. Morphologies are described in Supplemental Table S5.

(See separate PDF files for Figure S04 images)

**Figure S04. Exometabolomics of RCH2 wild-type and mutants cultured on N1B4 spent medium.** Isolate abundances not significantly different (ANOVA, Tukey HSD, p<0.05) are indicated with the same letter above their boxplots. In total, 125 metabolites were detected in the spent medium of cross-fed cultures and media controls. Eight metabolites were near completely depleted by all RCH2 WT and deletion mutant strains, including the purines adenine, guanine and hypoxanthine; the small organic acids glycerate, fumarate, glyceraldehyde/lactate, and malate; and the amino acid aspartate. An additional nine metabolites (2-amino-2-methylpropanoic acid, 2-oxovaleric acid, alanine, asparagine, glutamine, glutamic acid, glycine, malonate, pyruvate) were significantly decreased at varying amounts relative to each other. Forty metabolites were not significantly increased or decreased by any RCH2 WT and mutant strains. 2'-deoxyguanosine, citrulline, cytidine, dethiobiotin, guanosine, mandelic acid, thymidine and uridine were significantly increased by all RCH2 WT and mutant strains. Some metabolites were uniquely produced by one or more mutants relative to control medium and wild-type. In cases of reduced or negligible net utilization or increased amounts compared with the wild-type, these metabolites may be produced by the mutant or possibly released from cells due to lysis / stress (Fig. 4B).

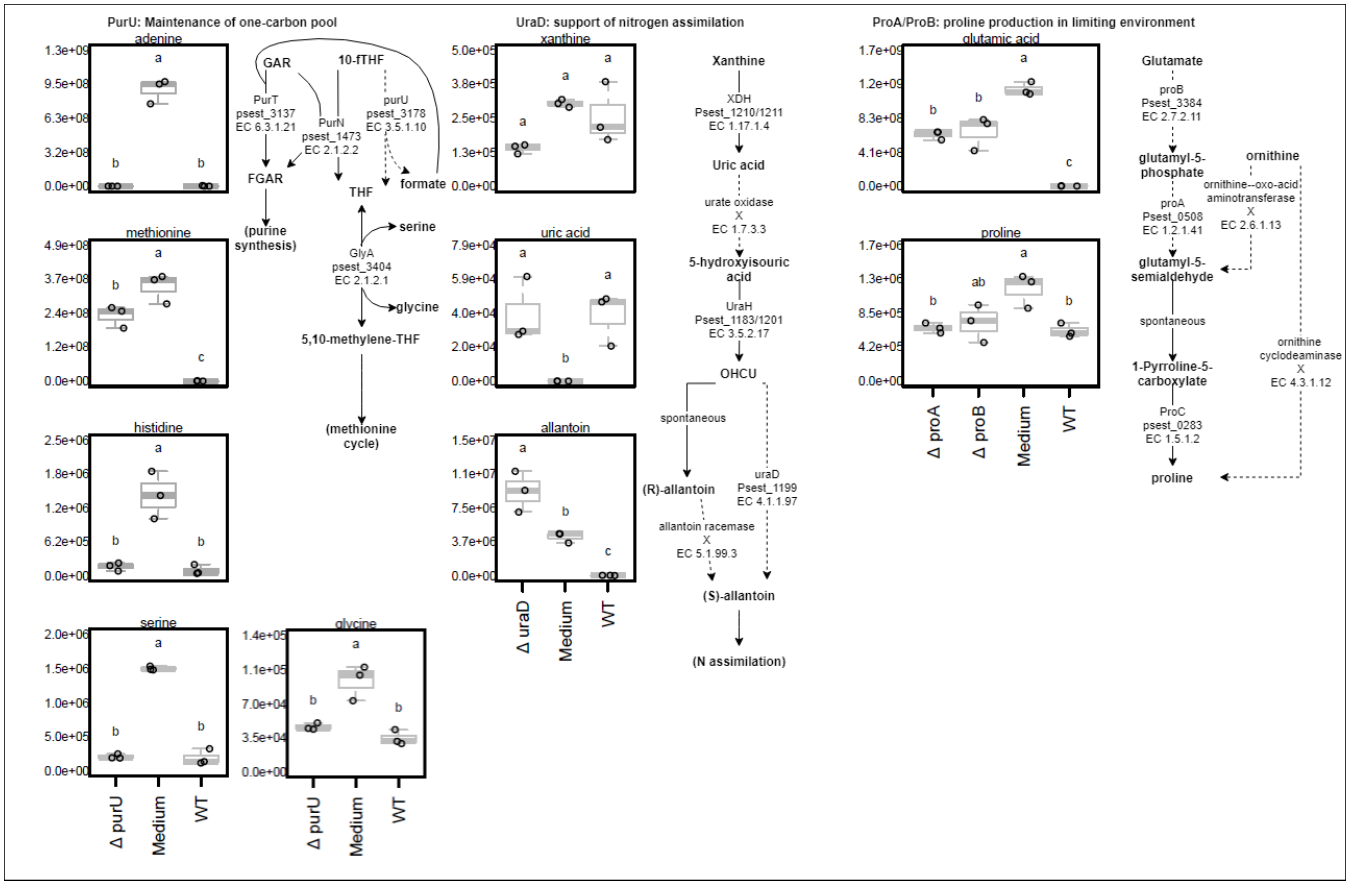


**Figure S05. Metabolomics of pathways important for RCH2 fitness on N1B4.** Anova/Tukey HSD results from each mutant compared with medium control and wild-type are displayed for metabolites associated with pathways contained mutated gene(s) of interest. Reactions (arrows) are annotated with enzymes when known along with gene ID and EC number; x indicates lack of known enzyme in RCH2. One-carbon metabolism: Formyltetrahydrofolate deformylase (PurU) converts 10-formyltetrahydrofolate to tetrahydrofolate and formate and plays a key role in maintaining formate pools and balancing the TFH:10f-THF ratio within the cell. PurU may be required in RCH2 for THF and formate production and stimulation of GlyA for maintenance of one carbon pool though folate and methionine cycles. In purine degradation, OHCU decarboxylase (UraD) converts 2-oxo-4-hydroxy-4-carboxy-5-ureidoimidazoline (OHCU) to S(+)-allantoin. Blastp search of amino acid sequences for urate oxidase from P. aeruginosa and Klebsiella pneumoniae resulted in no matches to a uricase in P. stutzeri RCH2, which would make it unable to use xanthine or uric acid for (S)-allantoin synthesis, but able to use 5-hydroxyisouric acid or OHCU (no standards available in library). Proline synthesis: Glutamate 5-kinase (ProB) and gamma-glutamyl phosphate reductase (ProA) sequentially catalyze the first two steps in the synthesis of proline. RCH2 appears to lack an ornithine cyclodeaminase (e.c 4.3.1.12) and Ornithine--oxo-acid aminotransferase (2.6.1.13) for alternative proline synthesis (mapping the protein sequence from P. stutzeri TS44 for 4.3.1.12 and from P. fluorescens for 2.6.1.13 resulted in no/very poor (<40%) matches in the NCBI database).
