## Supplementary figures and images for "Biofilm Interaction Mapping and Analysis (BIMA): A tool for deconstructing interspecific interactions in co-culture biofilms"

### SI Fig S04 pg1

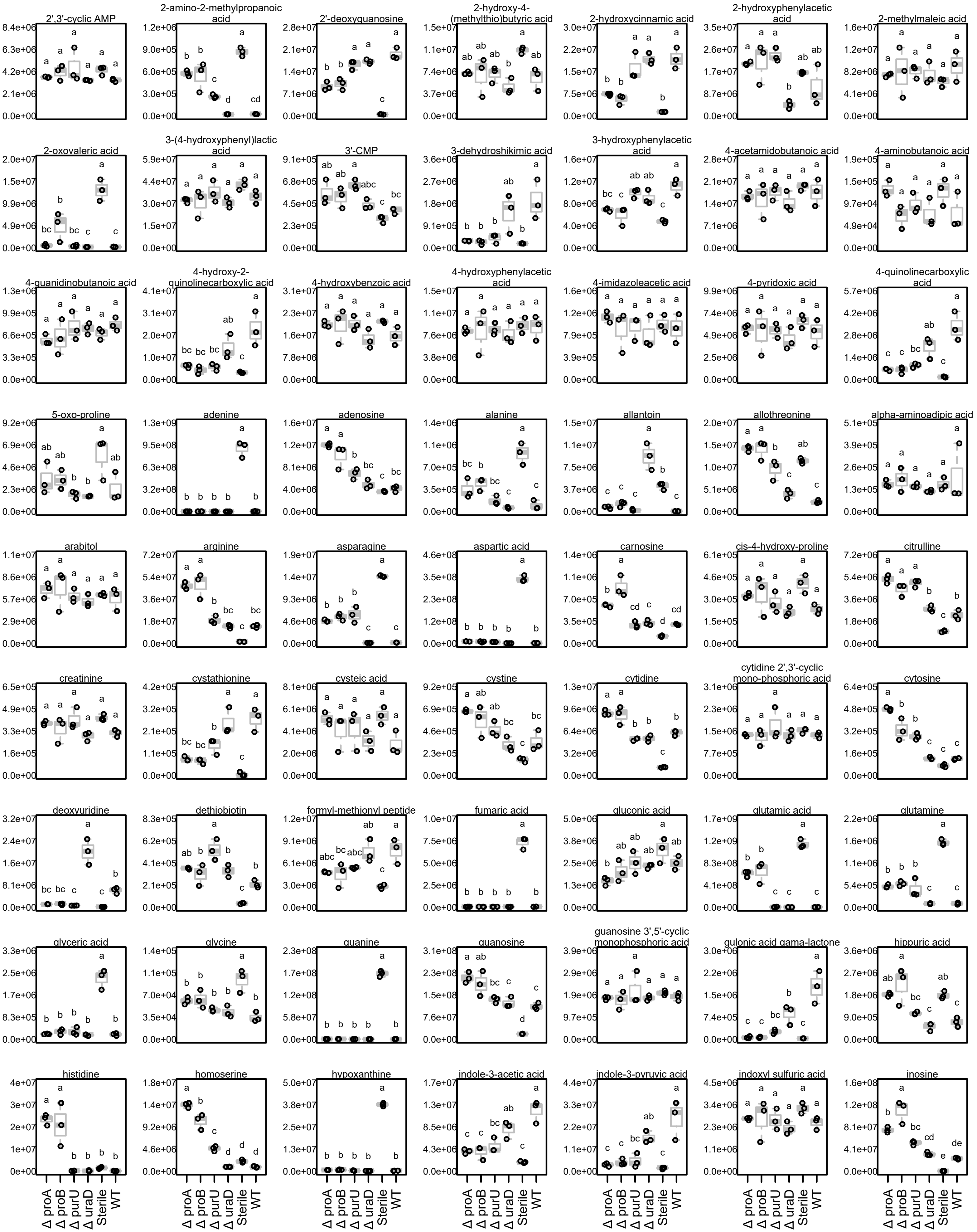

### SI Fig S04 pg2

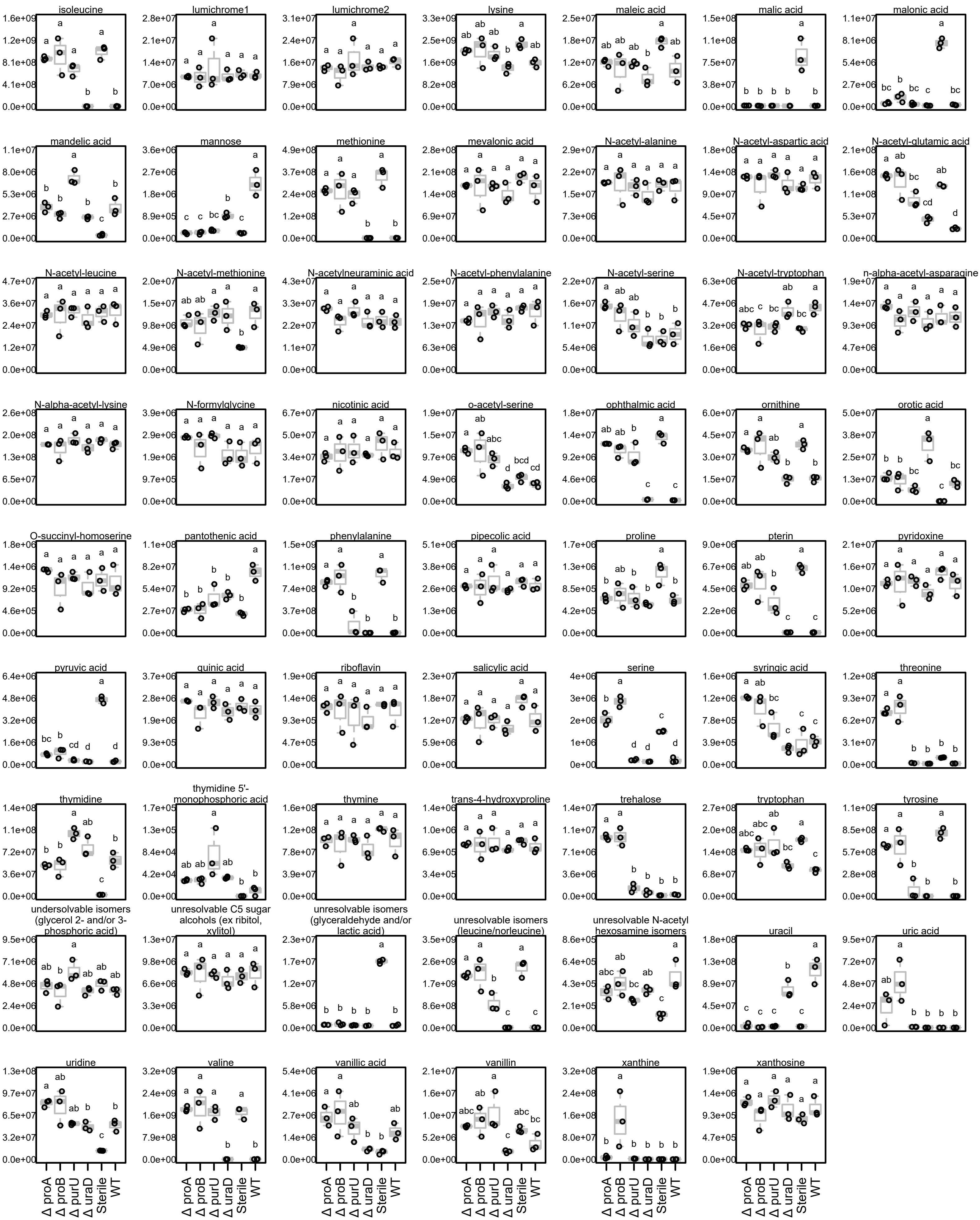
